# Evaluation of the adaptability of cassava genotypes in three different agro-ecological zones of Gabon

**DOI:** 10.1101/2024.06.05.597564

**Authors:** Branly Wilfrid Effa Effa, Rostand Abaga Moto, Yves Landry Zeh Nguema, Anaclé Ifada, Dick Sadler Demikoyo Kanghou, Stéphane Mibemu Guibinga, Sidoine Akoubou, Louis Clotaire Ngouang

## Abstract

Cassava is a staple food in over twenty African countries, including Gabon. However, current production cannot meet the needs of the Gabonese people. Studies are being carried out to find ways of increasing national production throughout the country. Our study of sixteen cassava genotypes planted in different agro-ecological zones of Gabon focuses on three quantitative variables: number of lobes per leaf, petiole length and plant height at the vegetative stage. This study was carried out to see which introduced cassava genotypes were adapted to environments with different rainfall levels.

## Introduction

Cassava (*Manihot esculenta* Crantz) is the staple food of 500 million in Africa, and its production is constantly increasing (Vernier et al., 2018). The interest in cassava is mainly focused on its tuberous roots’ rich in starch and its leaves rich in protein (Gnonlonfin et al., 2011). It is the fifth most important crop in the world behind maize, rice, wheat and potato (FAO, 2008), it also ranks second with production estimated at 32% of global production of tuberous roots and tubers (FAOSTAT, 2016). African continent produces annually 110 million tons of cassava roots, which makes Africa the world’s leading cassava producer followed by Asian continents with 55 million tons (Tetchi, 2007; Von Grebmer et al., 2013). Cassava is the staple food of many households in Central Africa, which explains the increasingly high levels of cassava consumption in line with population growth. In Central Africa, Gabon ranks 4th producer of cassava in 2017 (FAOSTAT, 2021). Cassava is eaten fresh or processed in producer countries (around twenty African countries, including Gabon) and provides around 50 kilo calories per person per day (Effa et al., 2024). It is also involved in the prevention of numerous diseases such as eye and cardiovascular diseases, and even cancers (Armstrong, 1999; Bauernfeind, 1981; Bast et al., 1998; Eisenreich et al., 2001). Its constituents, notably carotenoids, are thought to be involved in regulating the immune system and stabilizing the genome (Fraser and Bramley, 2004). However, due to abiotic and biotic constraints, current cassava production is unable to meet the needs of the local population (Cacaï et al., 2012). To deal with these various constraints, a number of strategies have been proposed, including the introduction of more resilient varieties. The aim of our study is to analyze the behavior of a dozen varieties outside their original environment, in the presence of local varieties.

## Materials and methods

### Plant material

This study was carried out on three sites with different rainfall rhythm, namely Makokou with an equatorial rhythm (1550 mm), Mouila with a transitional tropical rhythm (2000 mm), and Oyem with an equatorial rainfall rhythm (1750 mm) differing from that of Makokou in the quantities of annual rainfall received (Leonard & Richard, 1993). On each site, 16 cassava genotypes were planted with 50 individuals each.

### Experimental design

The experimental set-up used was complete randomization. The planting density was one meter between plants.

### Statistical analysis

The number of lobes per leaf and the length of the petiole are quantitative parameters that discriminate at the vegetative stage in cassava (Fukuda et al., 2010). The plant height parameter was considered to give us an idea of the trends that have emerged with regard to the adaptation of the introduced genotypes. The parameters number of lobes per leaf, petiole length and plant height were assessed for each genotype four months after sowing (at the vegetative stage). XLSTAT software was used for the analyses.

### Result

The results are presented for each site

## Number of lobes per leaf

At the Makokou site, the number of leaf lobes varies from 5.8 to 8 obtained respectively by genotype P0022 and genotype 960023 (Figure 1). Analysis of variance shows that there is a highly significant difference between genotypes for the parameter number of leaf lobes per leaf (*p-value* <0.0001).

**Figure 1:**
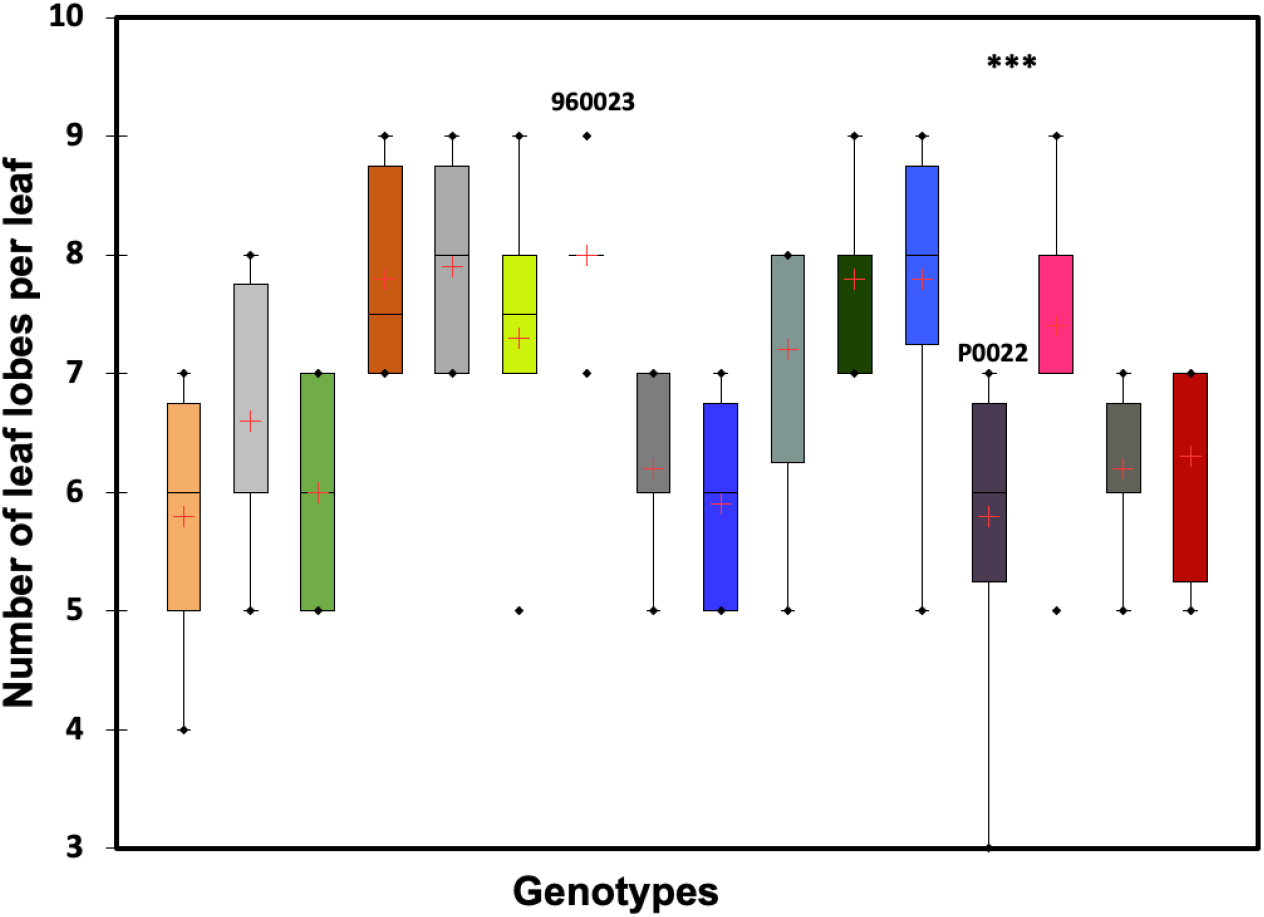
Number of leaf lobes per leaf and per genotype observed at Makokou, n=50. ***: *p-value* < 0.0001

At the Mouila site, the number of leaf lobes varies from 2.6 to 6.6 obtained respectively by genotype P0022 and genotype Moutoumbi (Figure 2). Analysis of variance shows that there is a highly significant difference between genotypes for the parameter number of leaf lobes per leaf (*p-value* < 0.0001).

**Figure 2:**
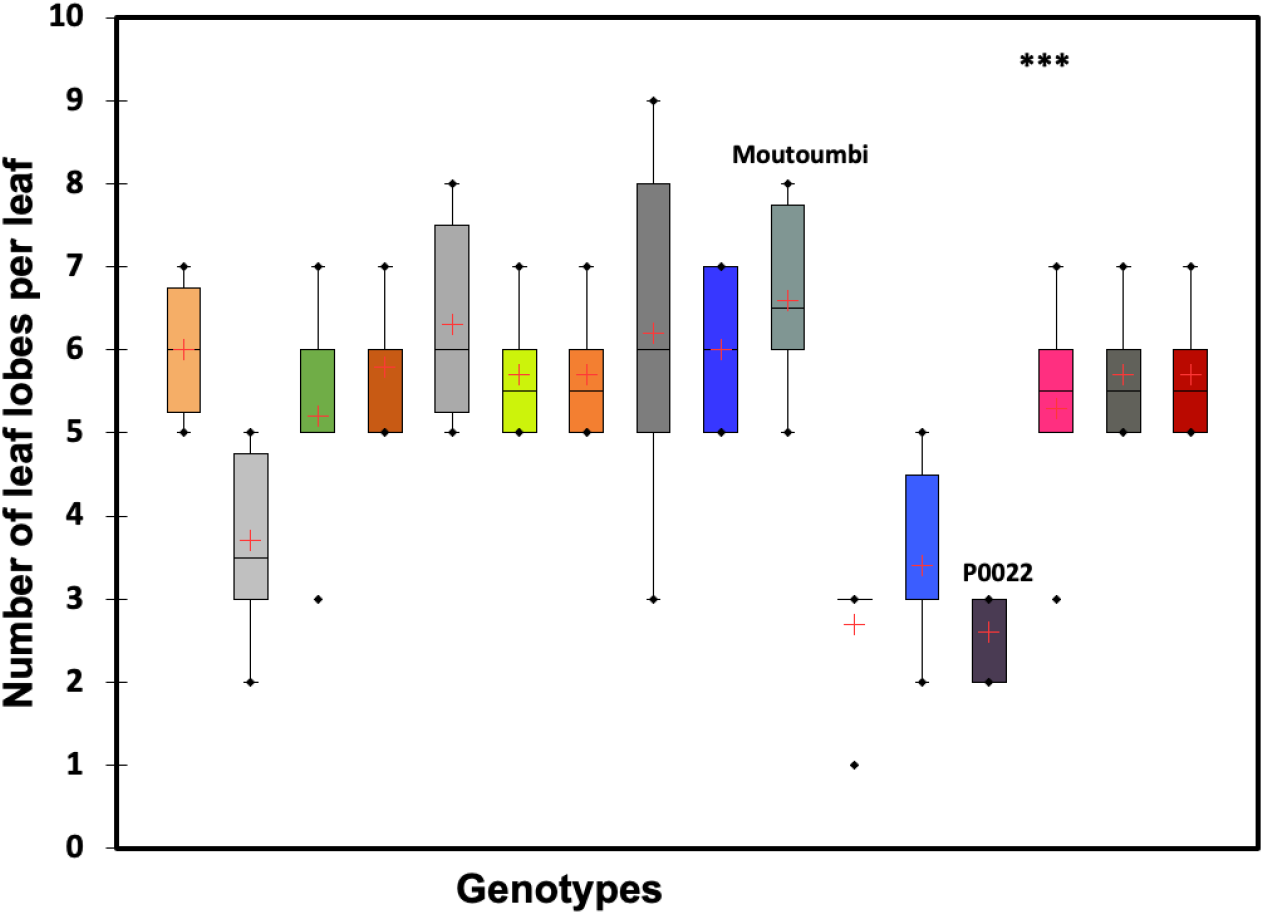
Number of leaf lobes per leaf and per genotype observed at Mouila, n=50. ***: *p-value* < 0.0001

At Oyem site, the number of leaf lobes varies from 4.1 to 7.8 obtained respectively by genotype 11797 and genotype Mambikini (Figure 3). Analysis of variance shows that there is a highly significant difference between genotypes for the parameter number of leaf lobes per leaf (*p-value* < 0.0001).

**Figure 3:**
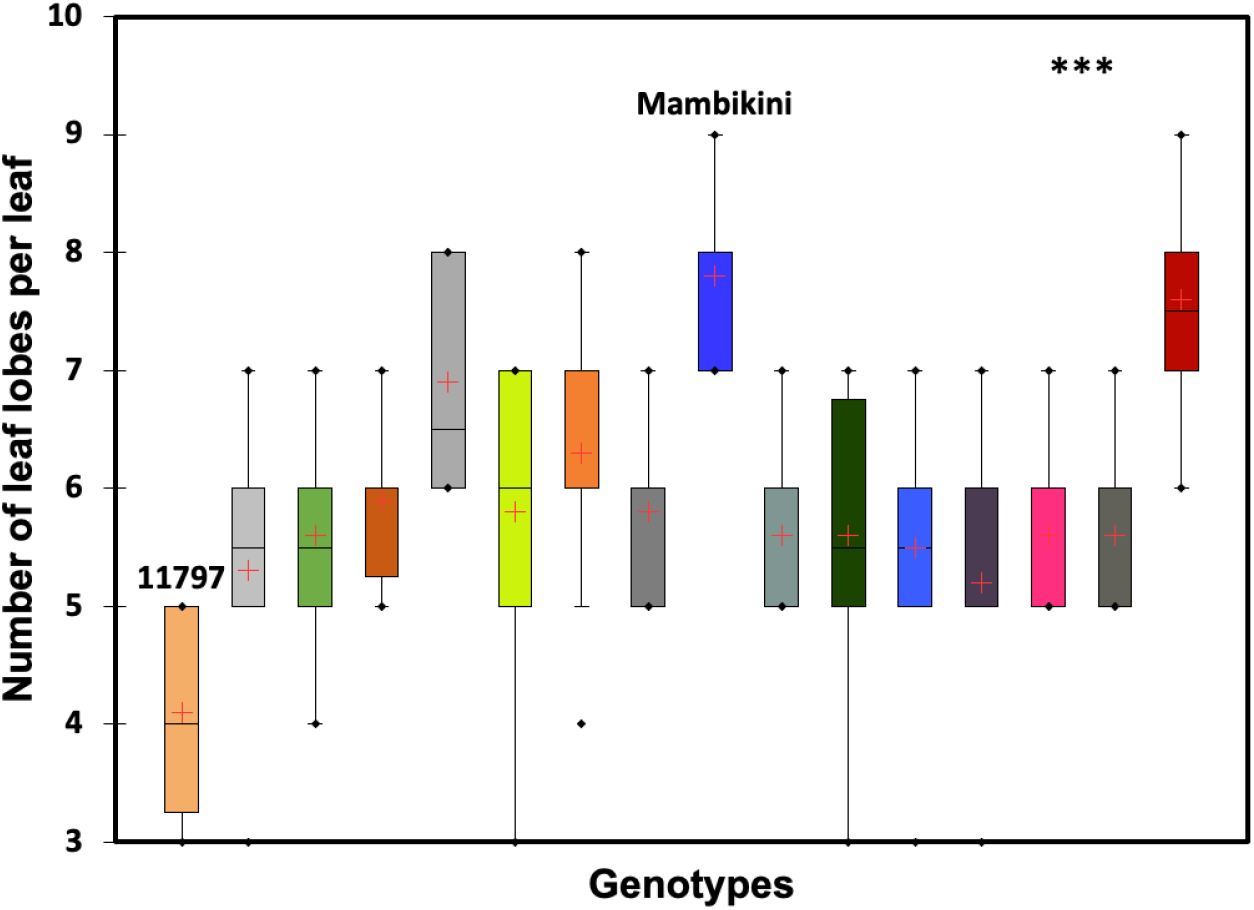
Number of leaf lobes per leaf and per genotype observed at Oyem, n=50. ***: *p-value* <0.0001

## Petiole length

At Makokou, the petiole length varies from 22.45 cm to 40.4 cm obtained respectively by genotype Banah and genotype Loka (Figure 4). Analysis of variance shows that there is a highly significant difference between genotypes for the parameter number of leaf lobes per leaf (*p-value* <0.0001).

**Figure 4:**
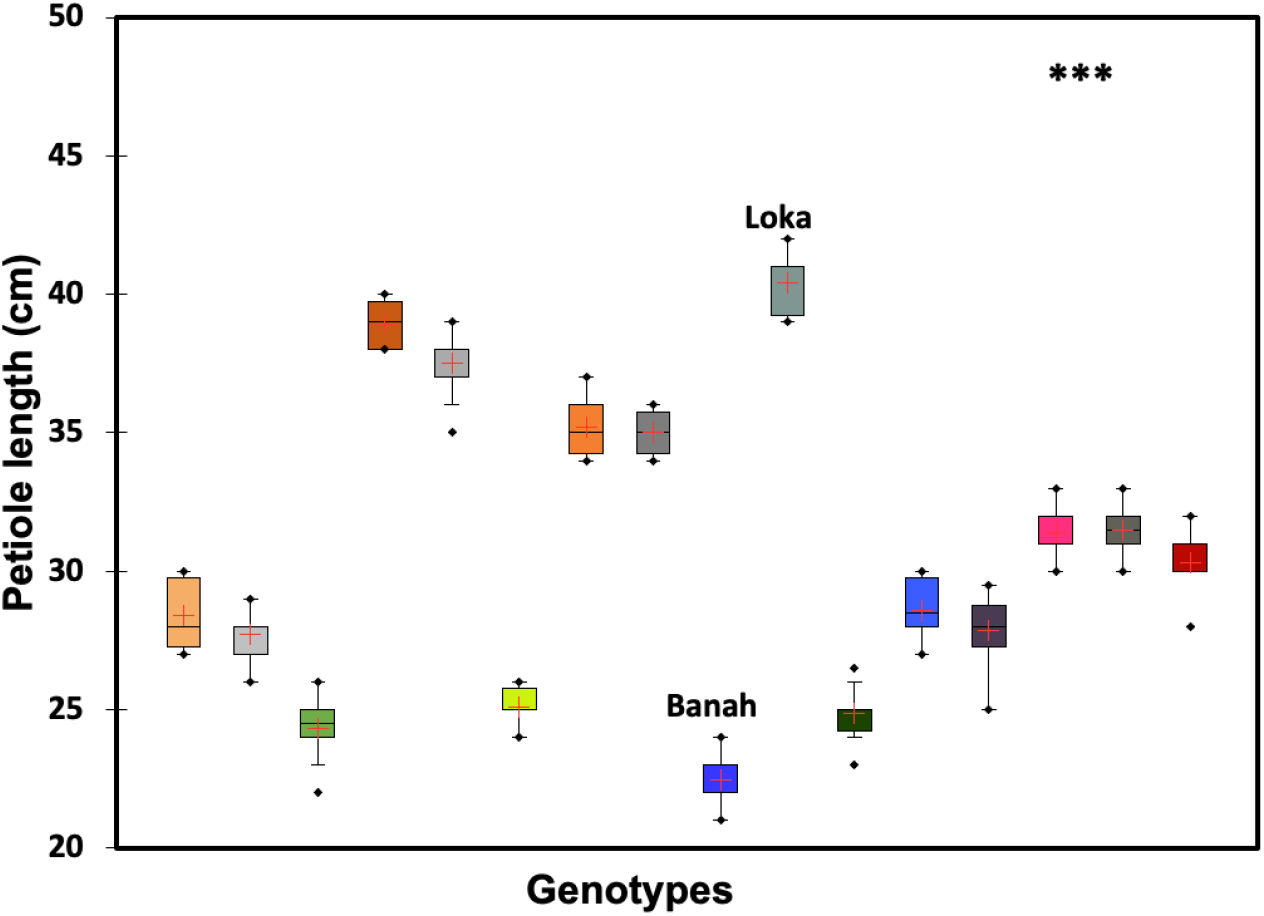
Petiole length per genotype observed at Makokou, n=50. ***: *p-value* < 0.0001

At Mouila, the petiole length varies from 9.2 cm to 30.65 cm obtained respectively by genotype P0044 and genotype 920326 (Figure 5). Analysis of variance shows that there is a highly significant difference between genotypes for the parameter number of leaf lobes per leaf (*p-value* < 0.0001).

**Figure 5:**
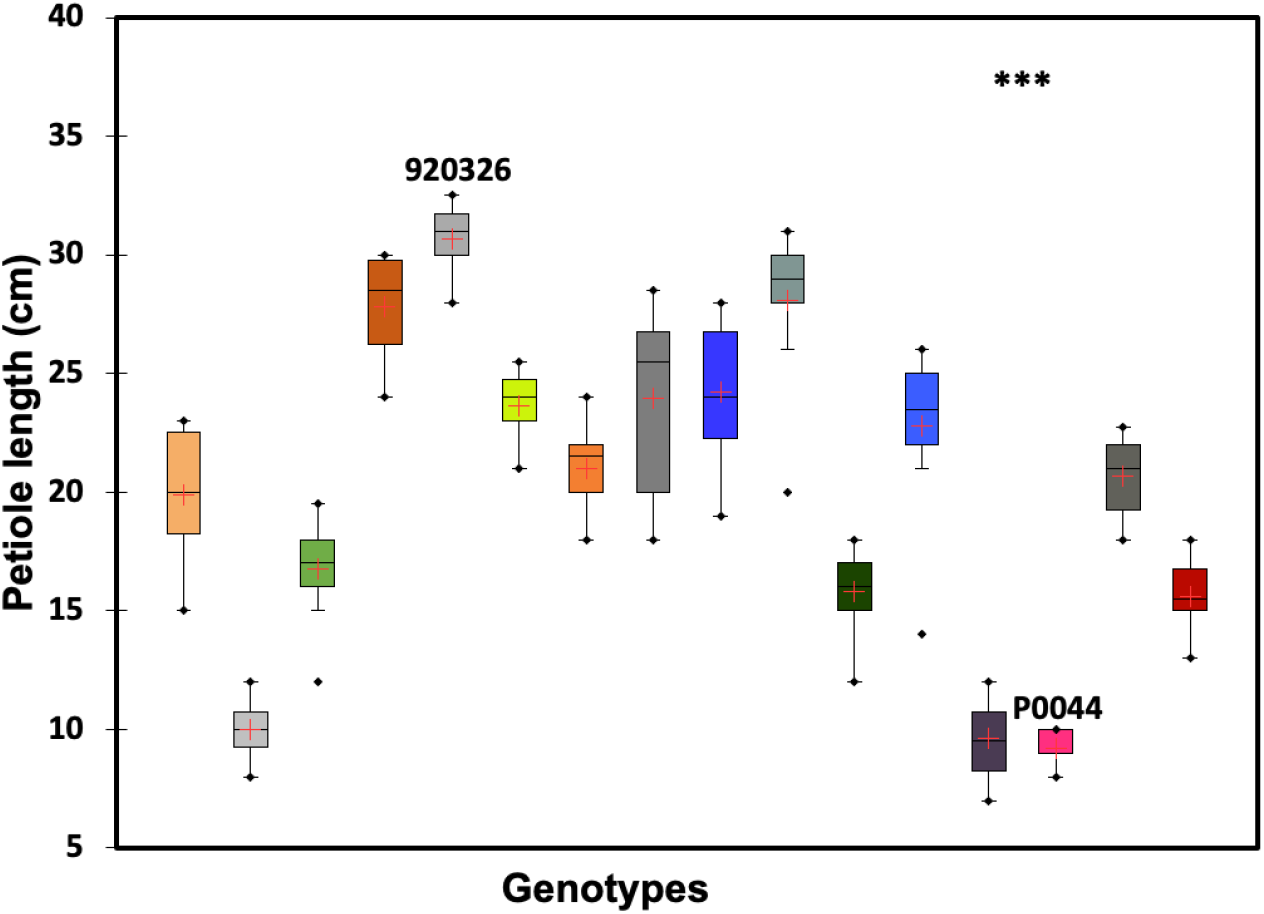
Petiole length per genotype observed at Mouila, n=50 and *p-value* < 0.0001

At Oyem, the petiole length varies from 15.25 cm to 34.1 cm obtained respectively by genotype P0003 and genotype Mambikini (Figure 6). Analysis of variance shows that there is a highly significant difference between genotypes for the parameter number of leaf lobes per leaf (*p-value* < 0.0001).

**Figure 6:**
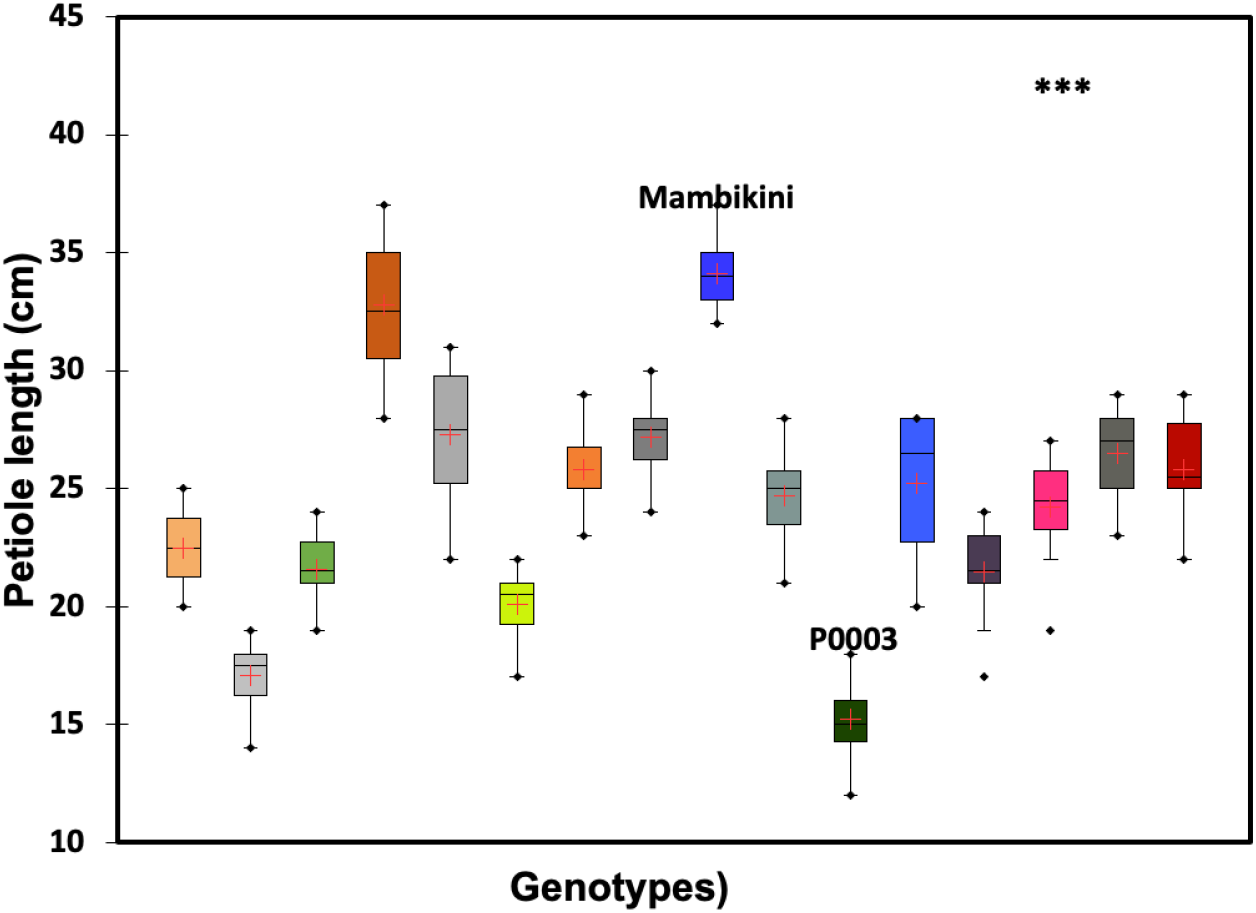
Petiole length per genotype observed at Oyem, n=50. ***: *p-value* < 0.0001

## Plant height

At Makokou, plant height varies from 165 cm to 348.2 cm obtained respectively by genotype P0003 and genotype 961414 (Figure 7). Analysis of variance shows that there is a highly significant difference between genotypes for the parameter number of leaf lobes per leaf (*p-value* <0.0001).

**Figure 7:**
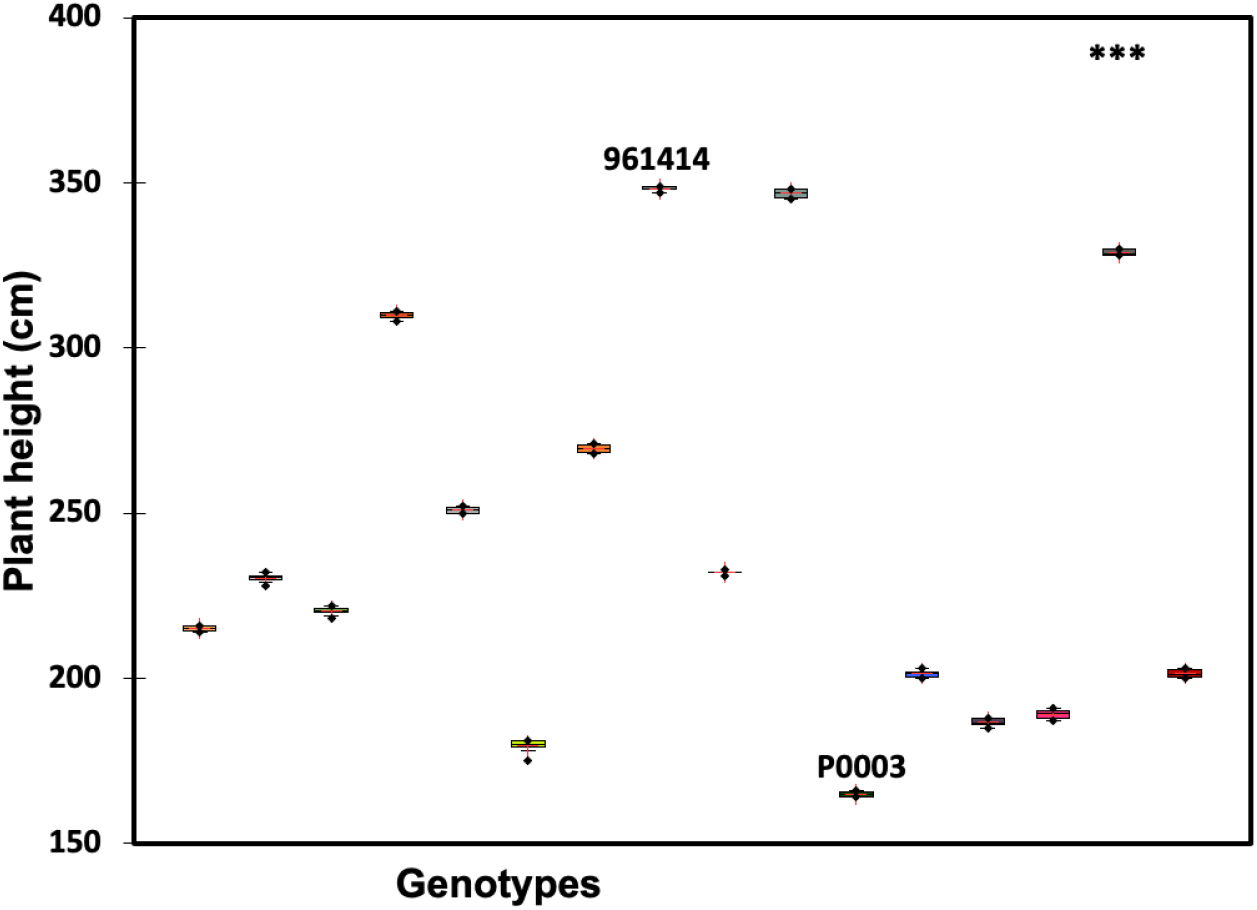
Plant height per genotype observed at Makokou, n=50. ***: *p-value* < 0.0001

At Mouila, plant height varies from 81.1 cm to 260 cm obtained respectively by genotype P0003 and genotype Ditadi (Figure 8). Analysis of variance shows that there is a highly significant difference between genotypes for the parameter number of leaf lobes per leaf (*p-value* < 0.0001).

**Figure 8:**
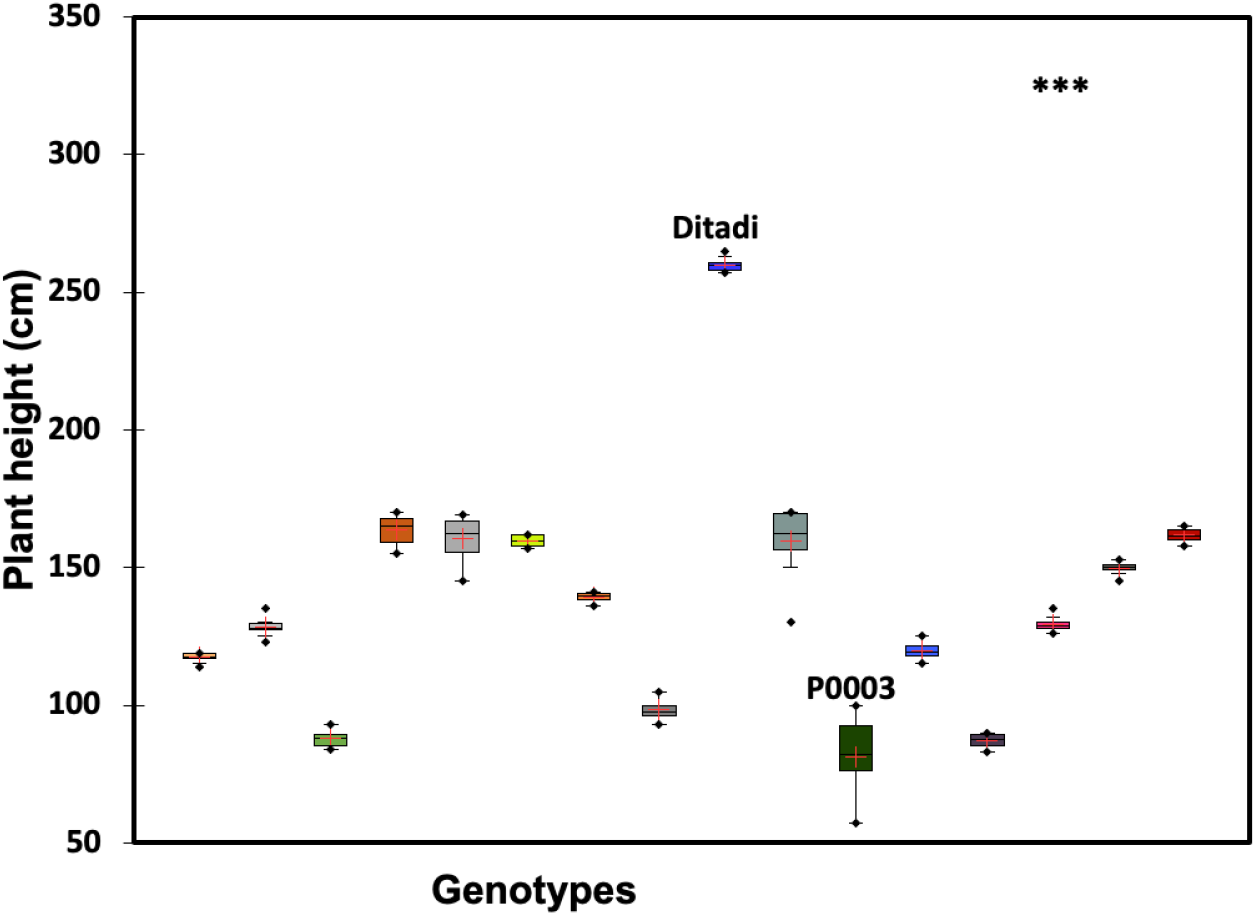
Plant height per genotype observed at Mouila, n=50. ***: *p-value* < 0.0001

At Oyem, plant height varies from 44.9 cm to 202.8 cm obtained respectively by genotype P0003 and genotype Mambikini (Figure 9). Analysis of variance shows that there is a highly significant difference between genotypes for the parameter number of leaf lobes per leaf (*p-value* < 0.0001).

**Figure 9:**
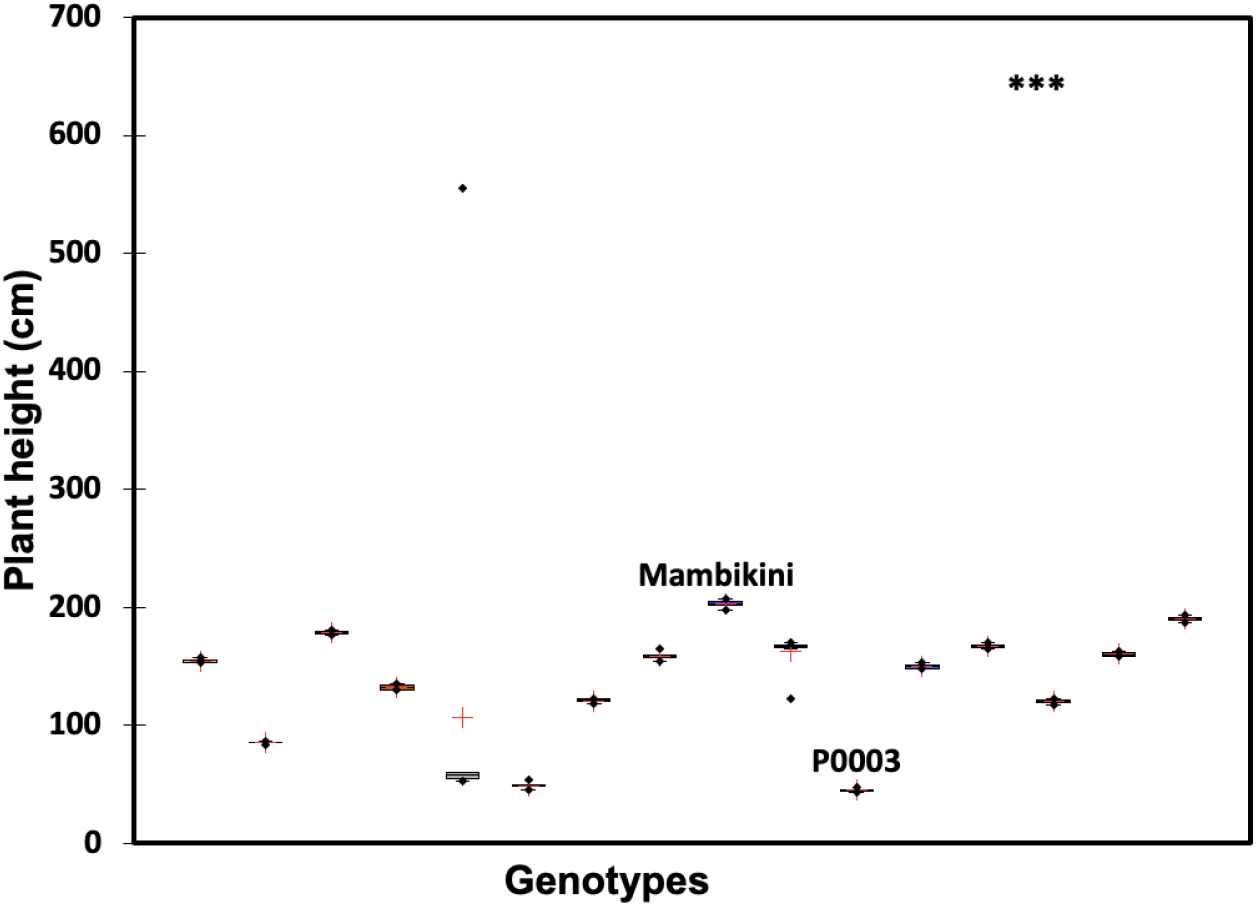
Plant height per genotype observed at Mouila, n=50. ***: *p-value* <0.0001

## Principal Component Analysis (PCA)

Principal Component Analysis carried out at the Makokou site with the quantitative variables number of leaf lobes per leaf, petiole length and plant size showed that genotype Loka was the best adapted to the environmental conditions (figure 10).

**Figure 10:**
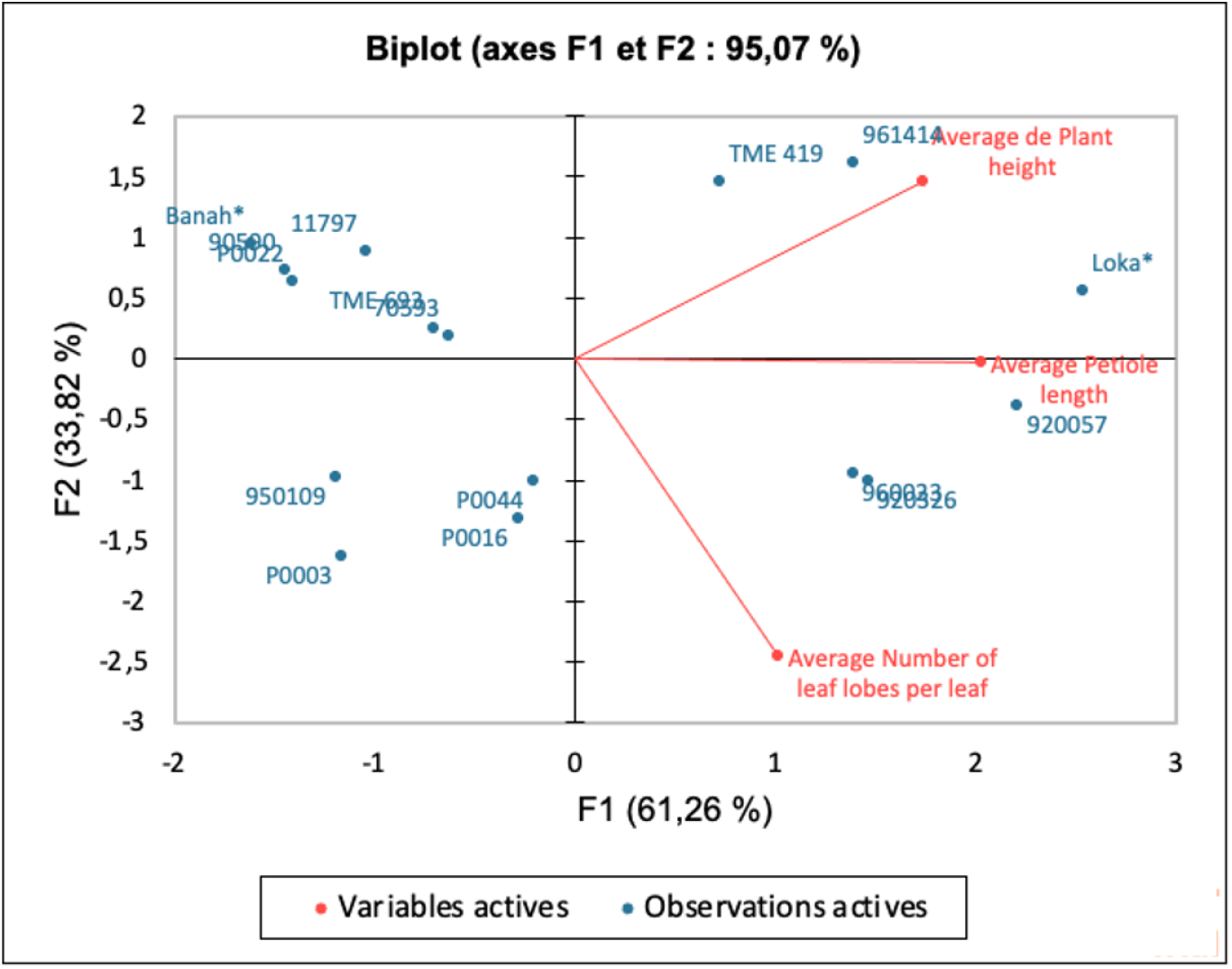
Principal Component Analysis of the Makokou site

Principal Component Analysis carried out at the Mouila site with the quantitative variables number of leaf lobes per leaf, petiole length and plant size showed that genotype Ditadi was the best adapted to the environmental conditions (figure 11).

**Figure 11:**
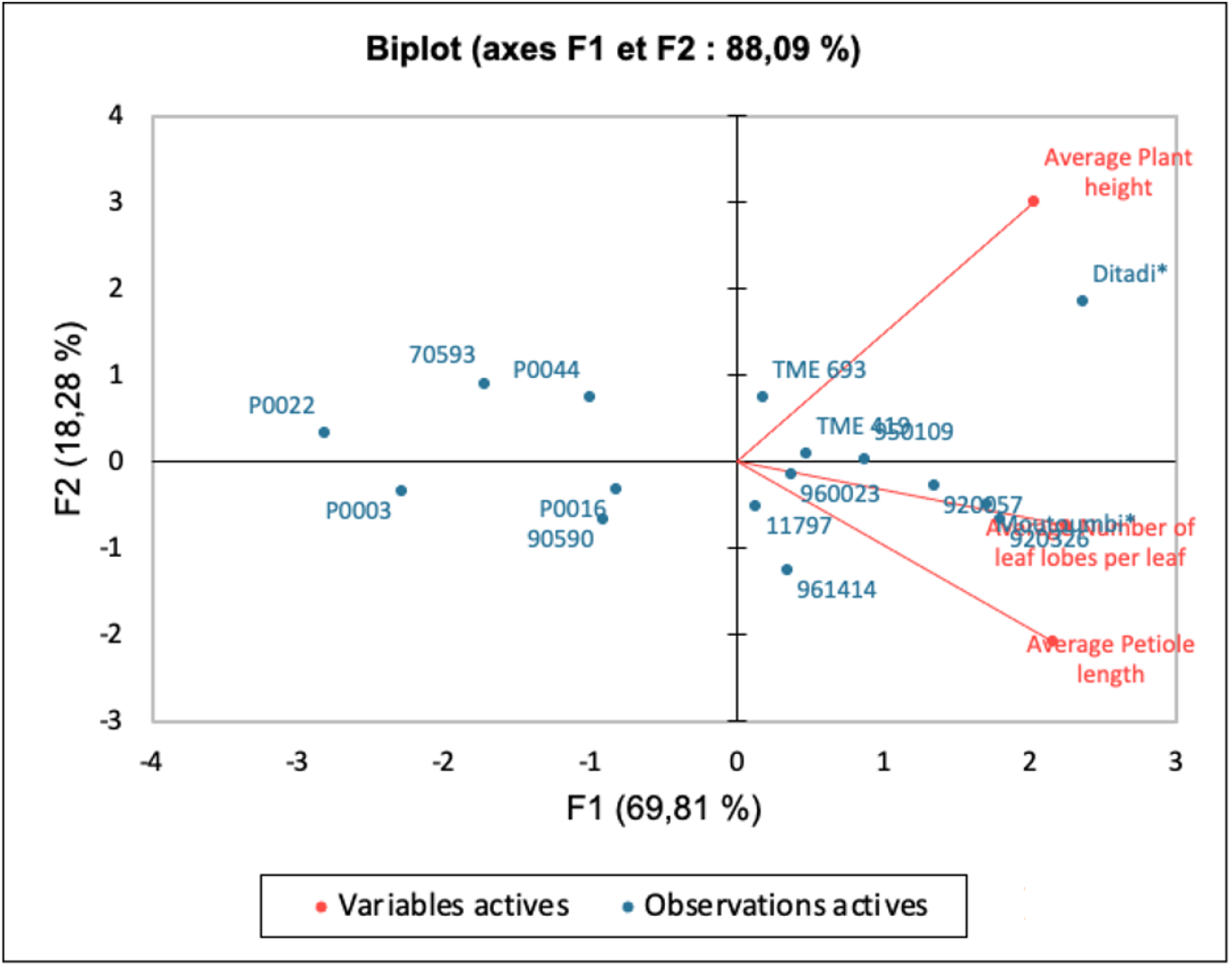
Principal Component Analysis of the Mouila site

Principal Component Analysis carried out at the Oyem site with the quantitative variables number of leaf lobes per leaf, petiole length and plant size showed that genotype Mambikini was the best adapted to the environmental conditions (figure 12).

**Figure 12:**
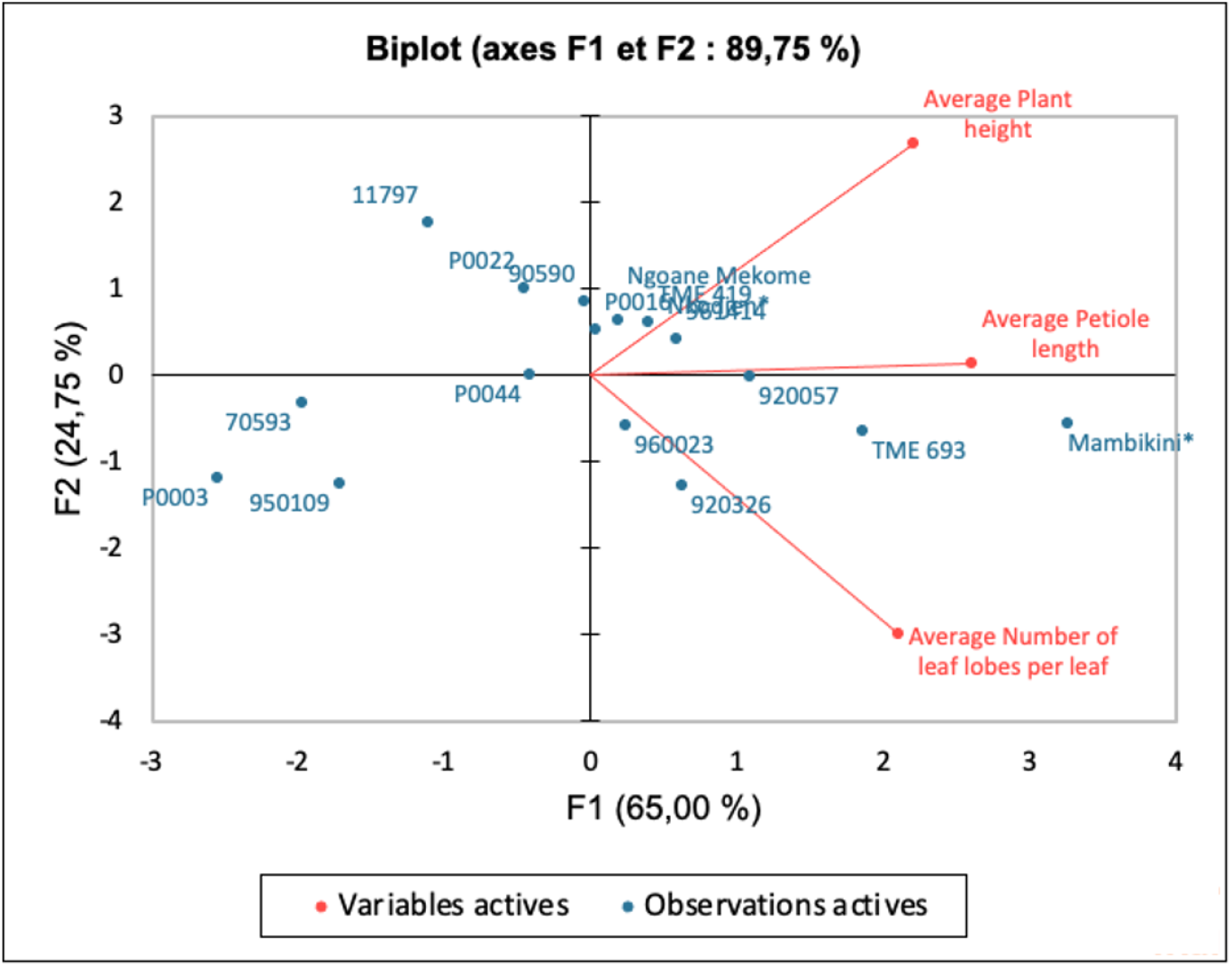
Principal Component Analysis of the Oyem site.

## Discussion

The experiment was carried out at each site with 14 genotypes introduced by the researchers and 2 local genotypes chosen by local growers. For each of the quantitative variables (number of leaf lobes per leaf, petiole length and plant height), the local genotypes had the highest values (in two of the three sites) compared with the introduced varieties. This confirms the choice of the populations for the use of these various local genotypes which were certainly selected because of their hardiness and their high tuberous root yields.

Principal Component Analysis revealed that local genotypes perform best at the vegetative stage. However, the results obtained after analyzing these quantitative variables at the vegetative stage merely give a trend, as tuberous root yields depend on other factors such as the amount of solar radiation captured by the leaves (Leepipatpaiboon et al., 2009; Mahakosee et al., 2022), which is also dependent on the amount of sunlight received by the leaves. Thus, changes including low levels of rainfall and its uneven distribution have a negative impact on the performance of a cassava genotype (Vincent et al., 2010; Sutrisno et al., 2023). This study focused on sites with different levels of annual rainfall because rainfall is a discriminating factor in assessing the performance of different cassava genotypes. The rainfall factor was also used to establish that local genotypes perform better than introduced genotypes at the vegetative stage using the quantitative variables number of leaf lobes per leaf, petiole length and plant height.

## Conclusion

The adaptation of plants to climate change is an important issue for maintaining or even increasing agricultural production. Our study, the results of which focused solely on the vegetative stage, shows that the most adapted genotypes at this stage are local genotypes. However, harvest results may differ due to other factors that influence tuberous root yields, including leaf area and root biomass, not to mention soil characteristics.

